## Supplementary Information for "HiExM: high-throughput expansion microscopy enables scalable super-resolution imaging"

**Table of Contents**

Fig. S1: Schematic depiction of HiExM device with displayed features.
Fig. S2: Overall schematic of HiExM comparing standard chemistry and photoinitiation.
Fig. S3: Detailed workflow of the HiExM protocol using photoinitiation.
Fig. S4: CF® conjugated antibodies yield robust signal and resistance to photobleaching in Irgacure HiExM.
Fig. S5: AcX and ProteinaseK titration.
Fig. S6: Residual hoechst signal occasionally remains underneath HiExM samples.
Fig. S7: Representative images post-expansion showing nanoscale resolution in HiExM.
Fig. S8: Infrequent higher expansion error in HiExM is due to abnormal stretching and tearing of the gel.
Fig. S9: Comparison of expansion error measurements between standard ExM chemistry and Irgacure 2959 photoinitiation.
Fig. S10: HiExM performs robustly on cells plated at low and high confluency.
Fig. S11: Nuclear volume/area analysis of Doxorubicin-treated cardiomyocytes before and after expansion.
Fig. S12: Nuclear edge analysis at different optical slices of Doxorubicin-treated cardiomyocytes.
Fig. S13: Line-scan analysis of Doxorubicin-treated cardiomyocytes.
Fig. S14: Comparison of derivative curves from Fig. 3 and Fig. S8.
Fig. S15: Nuclear edge analysis of $H_2O_2$-treated cardiomyocytes.
Fig. S16: Schematic depiction of the 24-well plate HiExM device.
Fig. S17: Example of a cell expanded in HiExM using Photo-ExM gel chemistry.

Movie S1. Example of gel expansion in APS/TEMED polymerized HiExM gels.
Movie S2. Example of gel expansion in Irgacure 2959 polymerized HiExM gels.
Movie S3. Visual representation of image analysis from Fig. 3.

Supplementary files: https://github.com/lboyerlab/hiExM_Supplementary_Files
Supplementary Methods
References

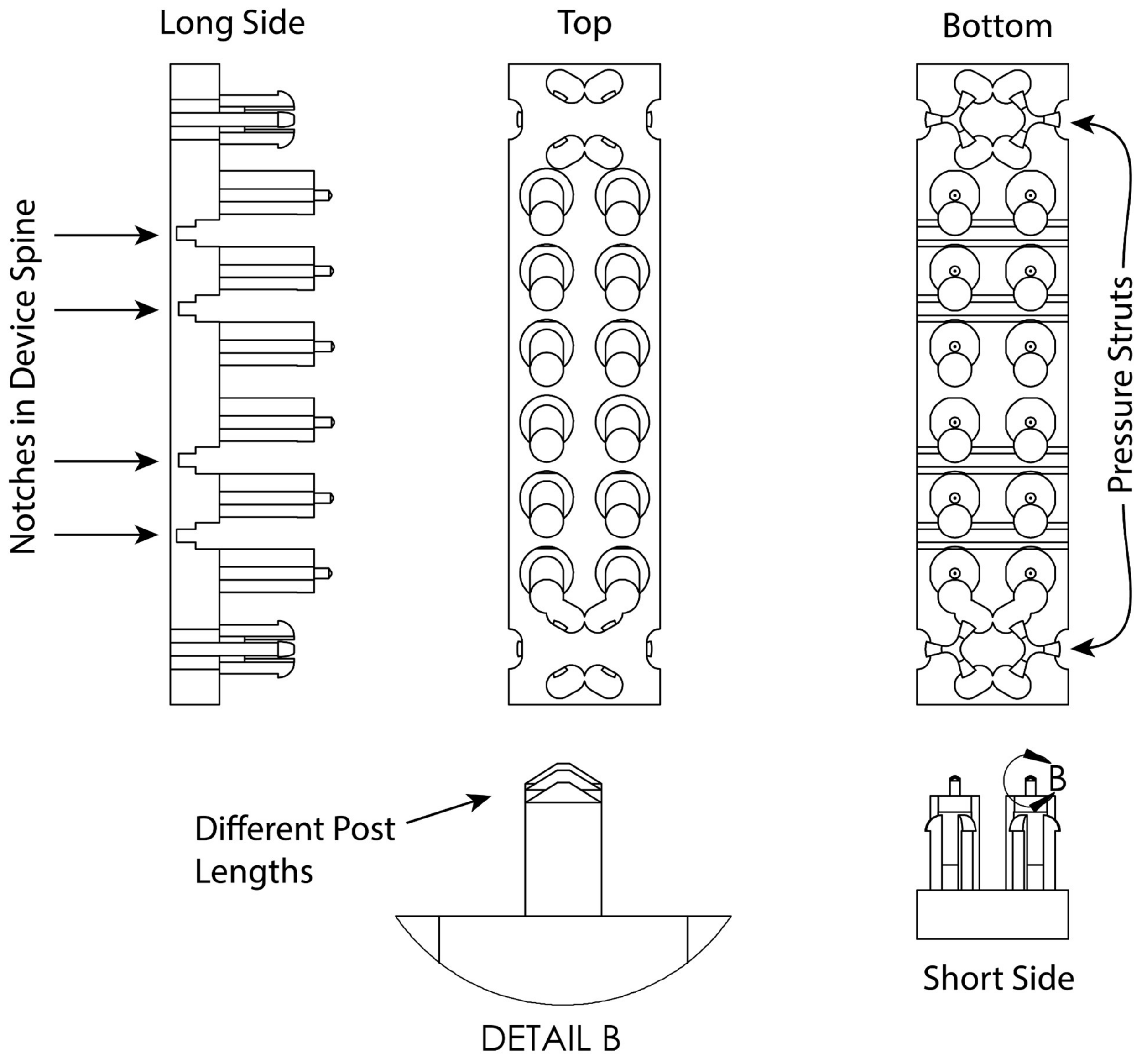


**Fig. S1: Schematic depiction of HiExM device with displayed features.** The device was designed with three different post lengths where the four center posts are the longest, the two pairs of posts on the ends near the pressure struts are the shortest, and the middle posts are an intermediate length. When the device is inserted into the well plate, the center four posts contact the culture surface first. As downward pressure is applied to the ends of the device (by the user pressing down above the pressure struts), the spine of the device bends at the notches such that all posts contact the plate. These design features allow for reproducible deposition of gel solution droplets and reproducible formation of toroidal gels within the well constraints.

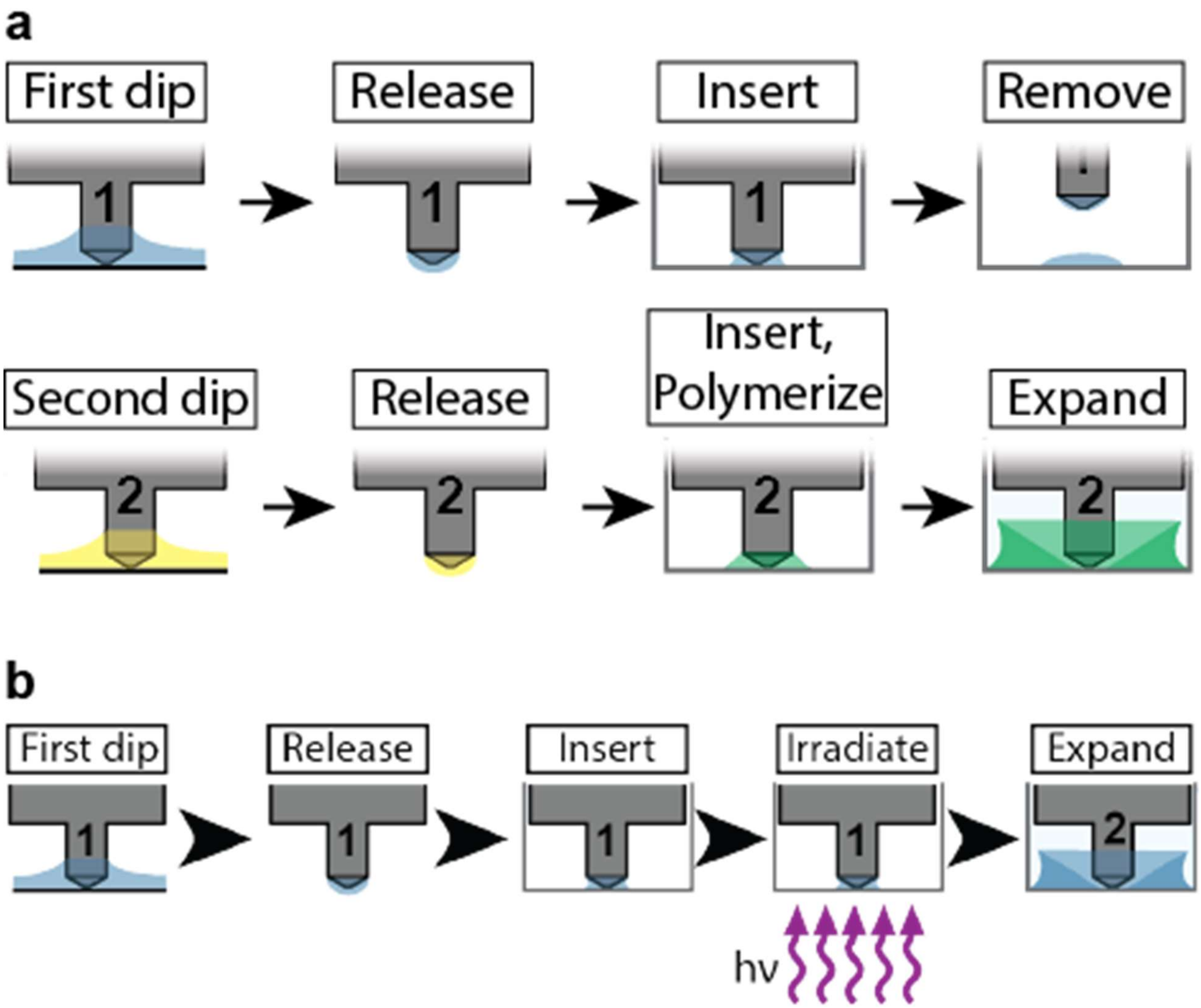


**Fig. S2: Overall schematic of HiExM comparing standard chemistry and photoinitiation.**
**A)** For standard ExM, the two initiating components (APS and TEMED) were added to cells in wells in a two-step process to control polymerization rate. In the first step, a droplet of TEMED containing gel solution is deposited in the well and the device is removed. In the second step, a second identical device delivers a droplet of APS containing gel solution to the same well with the TEMED containing droplet to initiate polymerization. The well plate is also kept on ice for 15 minutes while the APS and TEMED mixed, followed by heating the well plate at 50º C for 5 minutes to initiate rapid polymerization. Additionally, the protocol is performed in a glove bag purged twice with nitrogen. **B)** The use of photoinitiation HiExM as described in Methods allows for one step polymerization. The Irgacure 2959 gel solution is deposited and polymerization is initiated by irradiation with a UVA metal halide lamp.

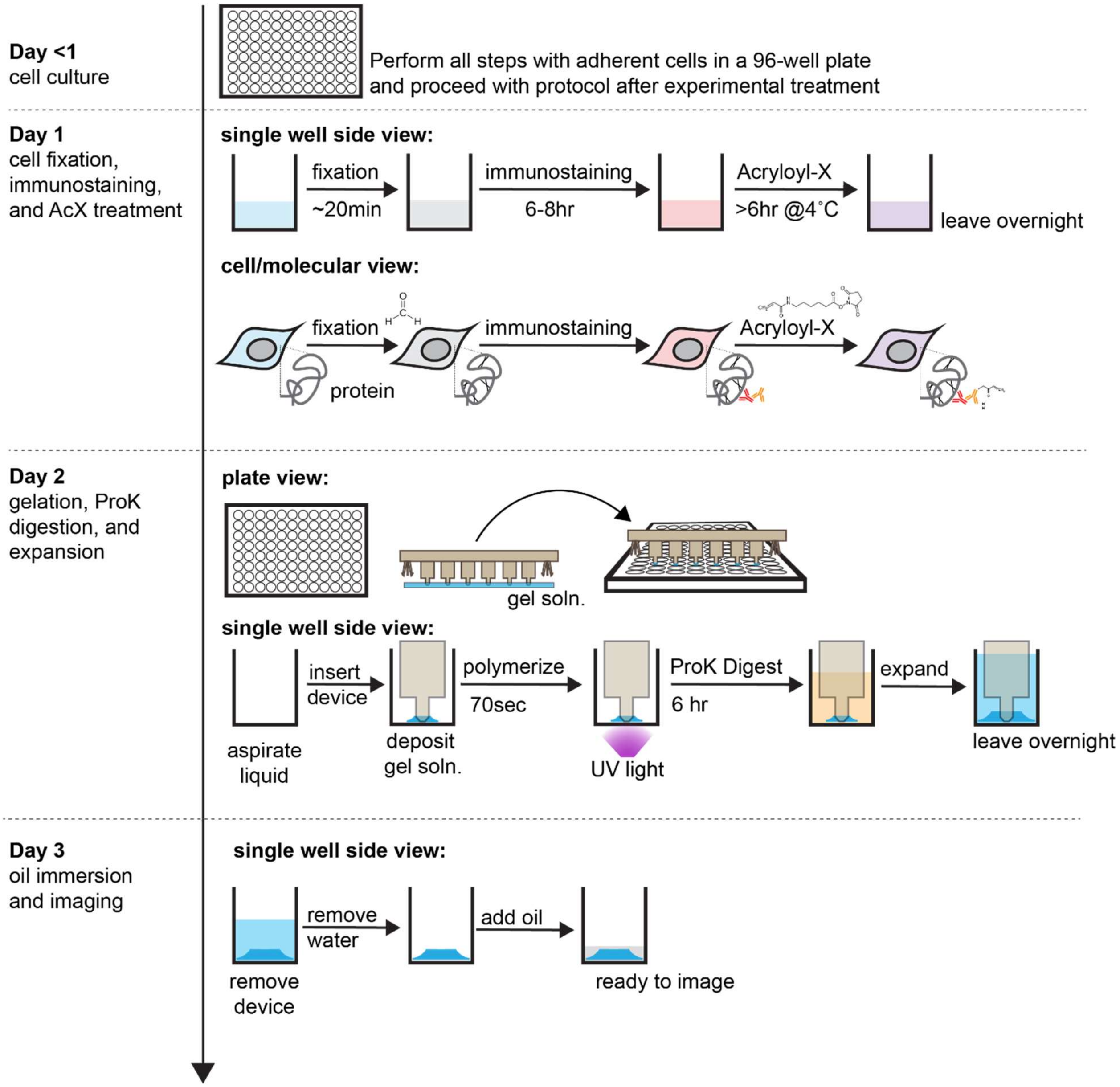


**Fig. S3: Detailed workflow of the HiExM protocol using photoinitiation.** HiExM can be used with cells cultured in commercially available 96-well plates. After conducting an experiment, cells can be fixed and stained as normal, followed by the addition of the anchor molecule Acryloyl-X. The following day, Acyloyl-X is aspirated, washed once with PBS and once with DI water, then aspirated again to leave cells dry. It is important to ensure that no residual liquid remains in the well. Following aspiration, the device is used to deposit gel droplets on cells. Deposited gels are left for ~90 seconds followed by a 70 second exposure to UV light to initiate polymerization. Proteinase K solution is then pipetted into the wells and after 6 hours, the plate is submerged in ~4L of water in a beaker and left suspended over a stir bar overnight. Water is then carefully aspirated from the wells and a layer of mineral oil is applied to each well to prevent evaporation and to stabilize the gel position for imaging. All steps in this procedure are performed at room temperature.

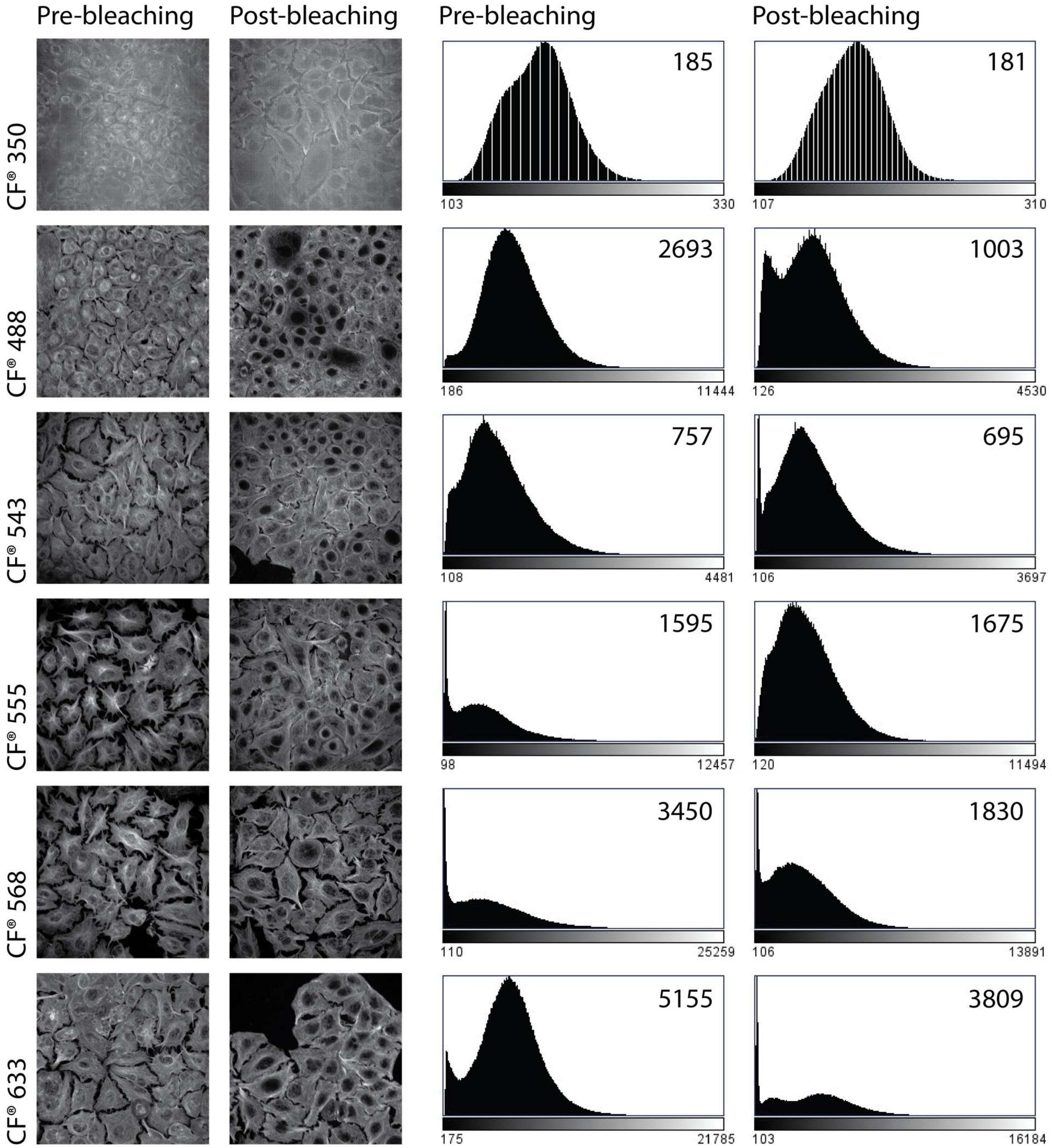


**Fig. S4: CF® conjugated antibodies yield robust signal and resistance to photobleaching in Irgacure HiExM.** Representative images of A549 cells stained with α-Tubulin antibodies and secondary antibodies conjugated to CF® dyes at different wavelengths before (left) and after (right) exposure to UV light as described in the HiExM protocol. Stained cells were exposed to UV light in the presence of Irgacure 2959 in 0.1% PBS. Histograms representing the raw pixel values from these images are shown on the right, where the number on each plot represents the peak pixel value for that image. Decreased peak pixel values indicate a loss of signal due to photobleaching.

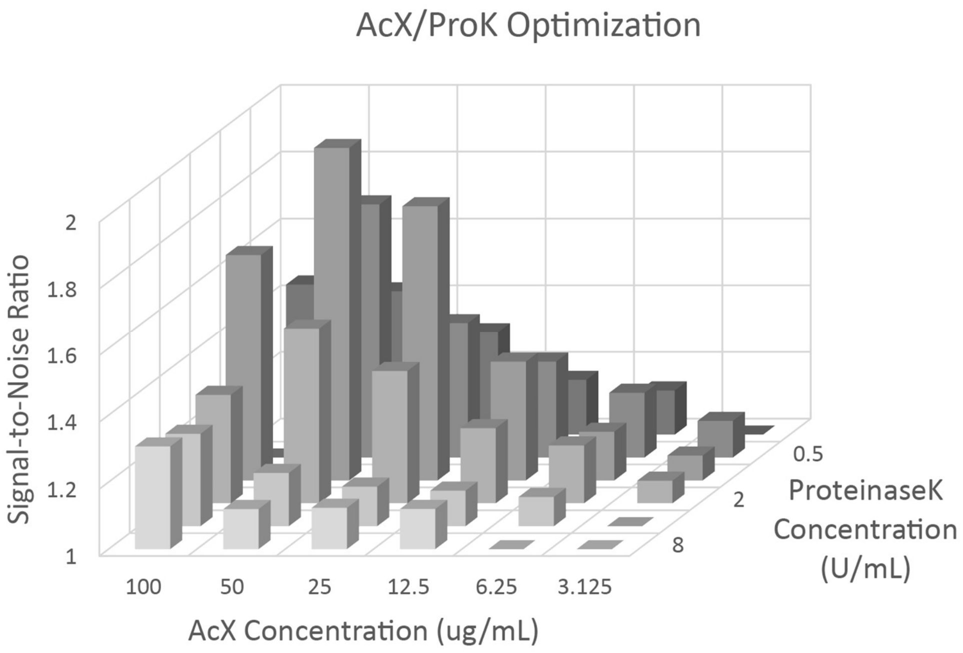


**Fig. S5: AcX and ProteinaseK titration.** Signal-to-noise ratio was determined by measuring the average fluorescence intensity in an area of the cell and an area of the background for α-tubulin stained A549 cells from images obtained using the Opera Phenix. Optimum signal-to-noise was found at concentrations of 50 μg/mL AcX and 1 U/mL ProteinaseK.

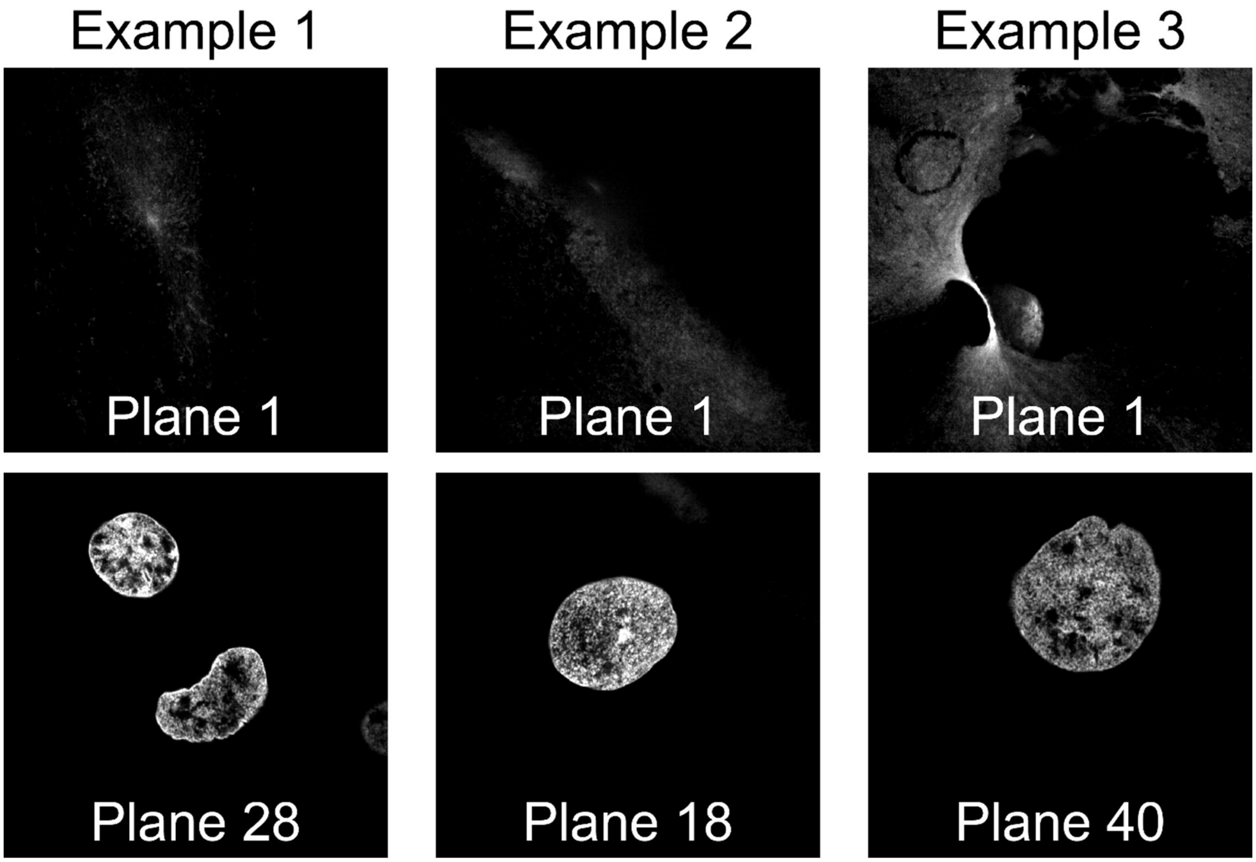


**Fig. S6: Residual hoechst signal occasionally remains underneath HiExM samples.** The lowest image planes (top row) of three example images that show residual hoechst signal. Higher planes from the same images (bottom row) show nuclei within the sample.

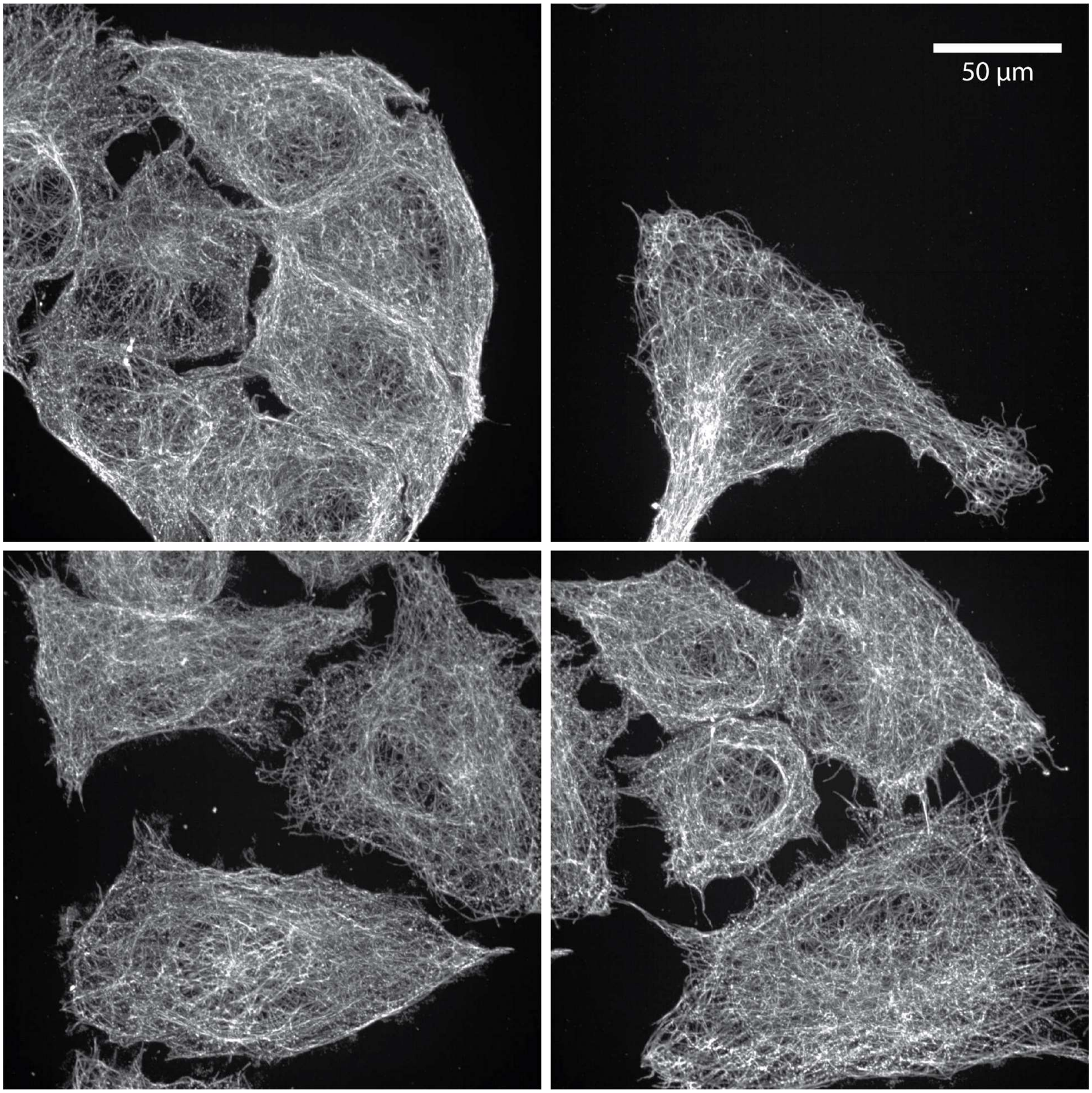


**Fig. S7: Representative images post-expansion showing nanoscale resolution in HiExM.** A549 cells were stained for α-Tubulin and imaged on an Opera Phenix High Content Screening System (PerkinElmer) using a 63x objective (1.15 na) as described in Fig. 2. Scale bar represents the real scale of the images, not corrected for expansion.

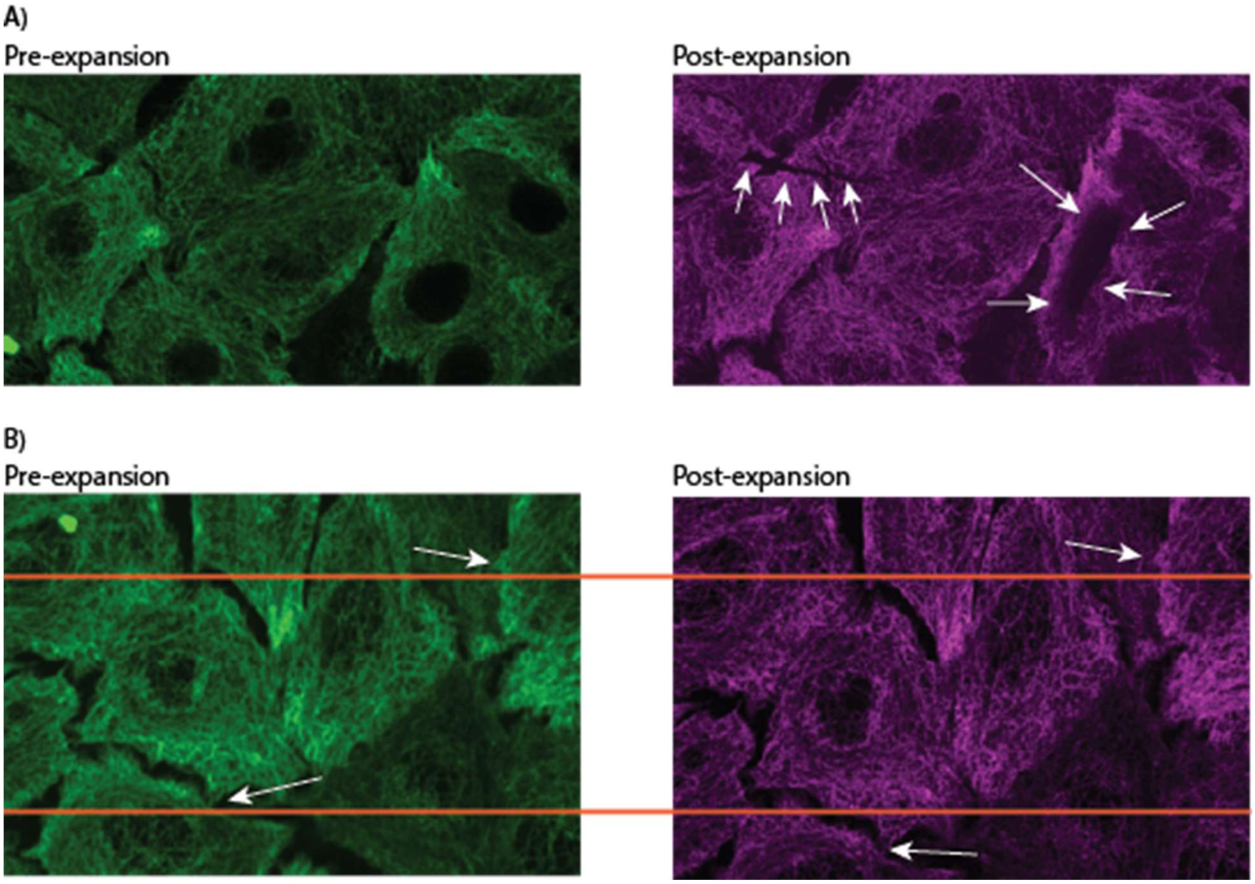


**Fig. S8: Infrequent higher expansion error in HiExM is due to abnormal stretching and tearing of the gel. A)** Arrows in the image on the right point to occasional artifacts arising during the expansion process. These rare artifacts generally occur closer to the center of the gel. **B)** Arrows point to matched features in the pre-expansion and post-expansion images. These images were aligned using TurboReg as previously described[1]. A comparison of the two images shows that the expanded sample was stretched anisotropically over length scales on the order of the size of the cell.

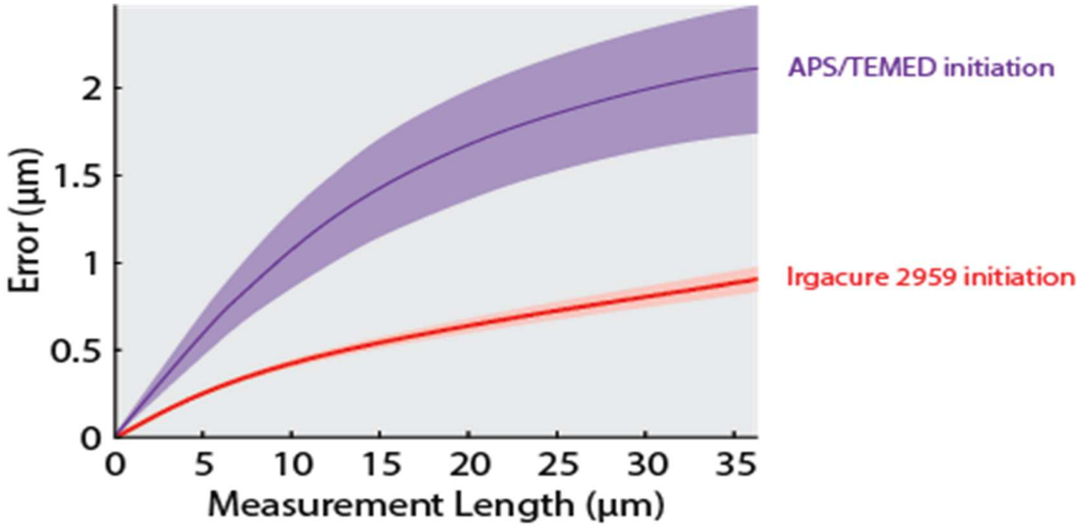


**Fig. S9: Comparison of expansion error measurements between standard ExM chemistry and Irgacure 2959 photoinitiation.** Consensus error curves were generated as described in Fig. 2 for APS/TEMED gels (left) as compared to Irgacure 2959 gels (right). These data demonstrate that photoinitiation results in robust expansion and less error compared to the APS/TEMED chemistry in our HiExM method.

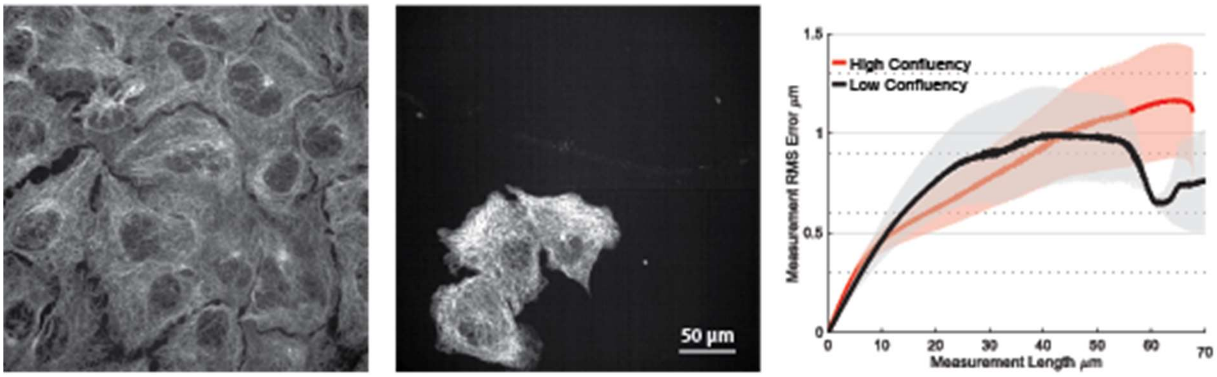


**Fig. S10: HiExM performs robustly on cells plated at low and high confluency.** Example images of high (left) and low (center) confluent cultures of A549s. Consensus error curves (right) were calculated as described in Fig. 2, show minimal error within ~0.2 um (n=4 fields of view for each condition).

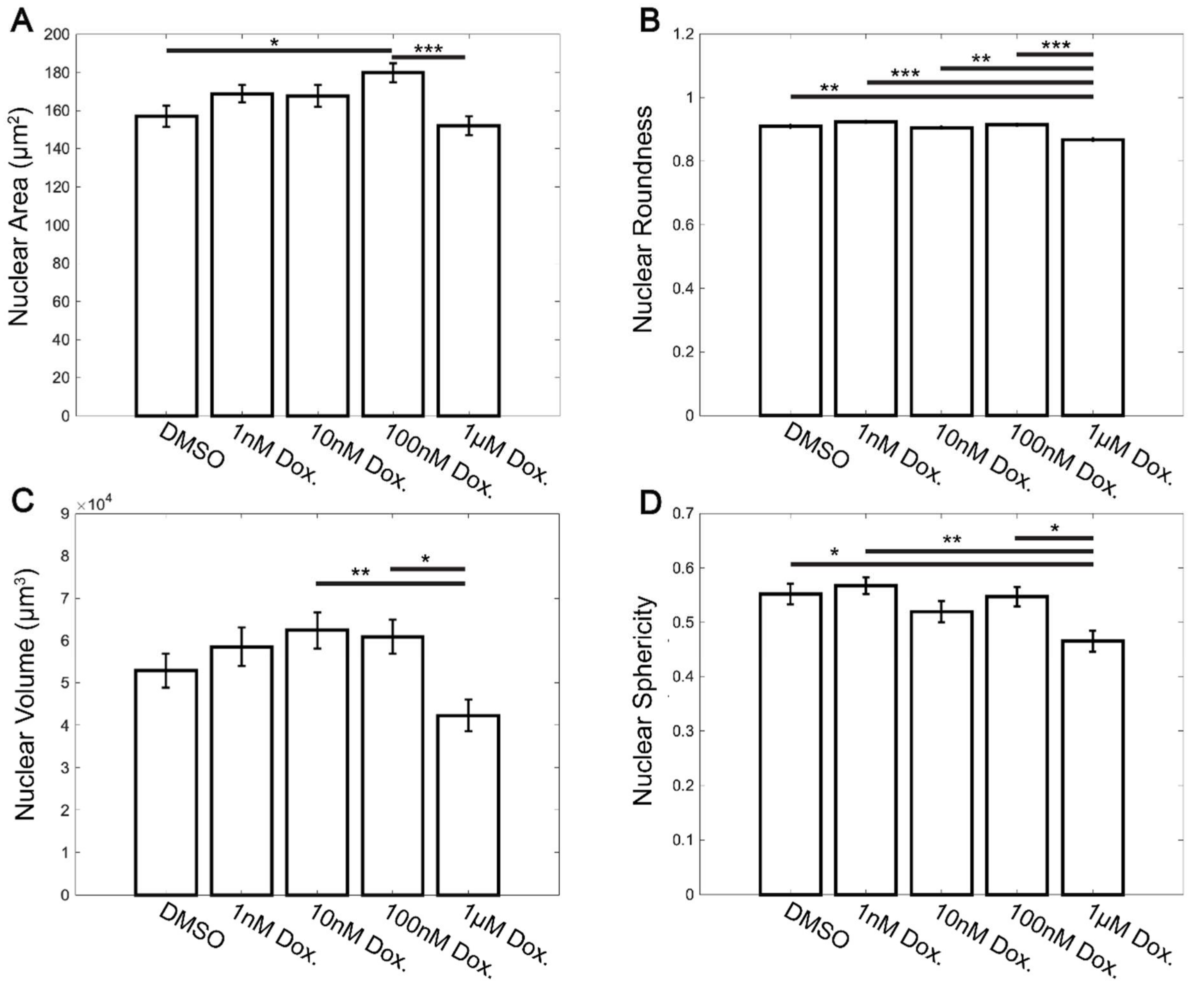


**Fig. S11: Nuclear volume/area analysis of Doxorubicin-treated cardiomyocytes before and after expansion.** Nuclear area (top left) and roundness (top right) were measured in pre-expanded cardiomyocytes. Nuclear volume (bottom left) and sphericity (bottom right) were measurable in expanded cardiomyocytes due to the increased axial resolution of HiExM. These measurements were performed in Harmony (Perkin Elmer). Error bars represent SEM and sample sizes are the same as in Fig. 3 for both unexpanded and expanded cells. * denotes significance from an ANOVA ( *: $p<0.05$, **: $p<0.005$, ***: $p<0.0005$).

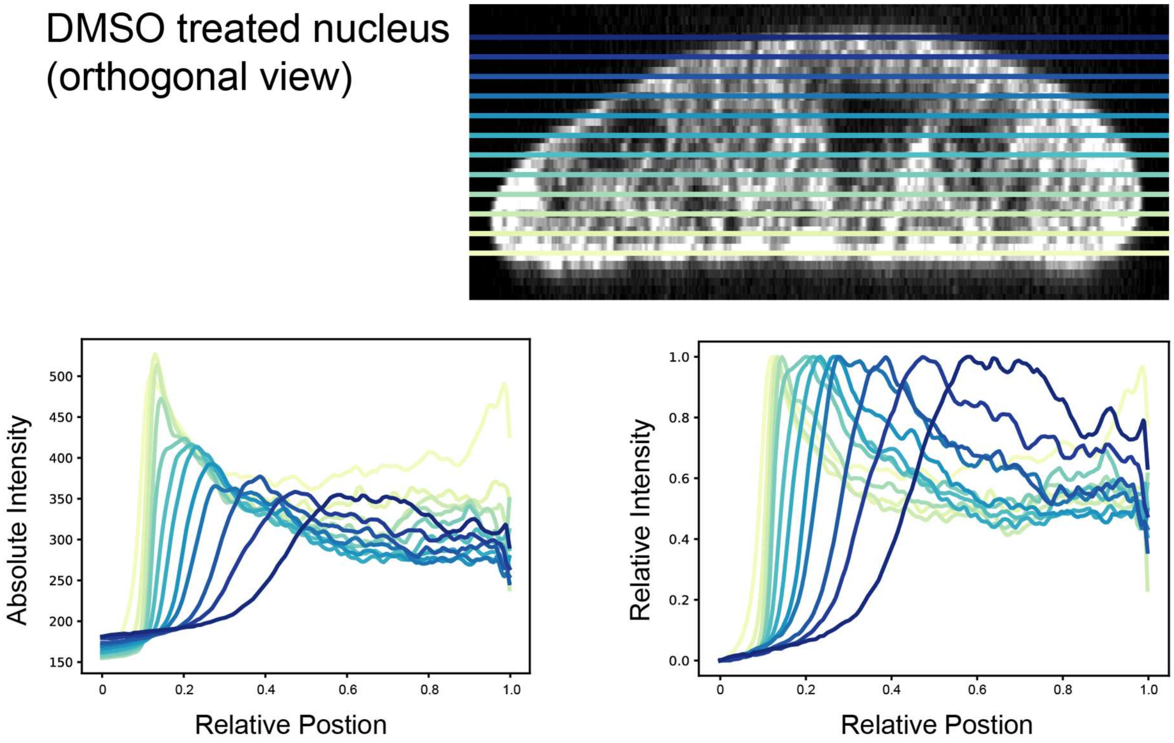


**Fig. S12: Nuclear edge analysis at different optical slices of Doxorubicin-treated cardiomyocytes.** Defining the optical plane that intersects the nucleus at a right angle in the z-axis is necessary for nuclear edge analysis. Orthogonal view of a DMSO-treated nucleus. Colored lines represent optical slices corresponding to the analysis results below the image (top). Absolute signal intensity as a function of relative position at 12 different z-slices using the 'ring-scan' analysis (bottom left). Relative signal intensity as a function of relative position at the same 12 z-slices using the same analysis (bottom right).

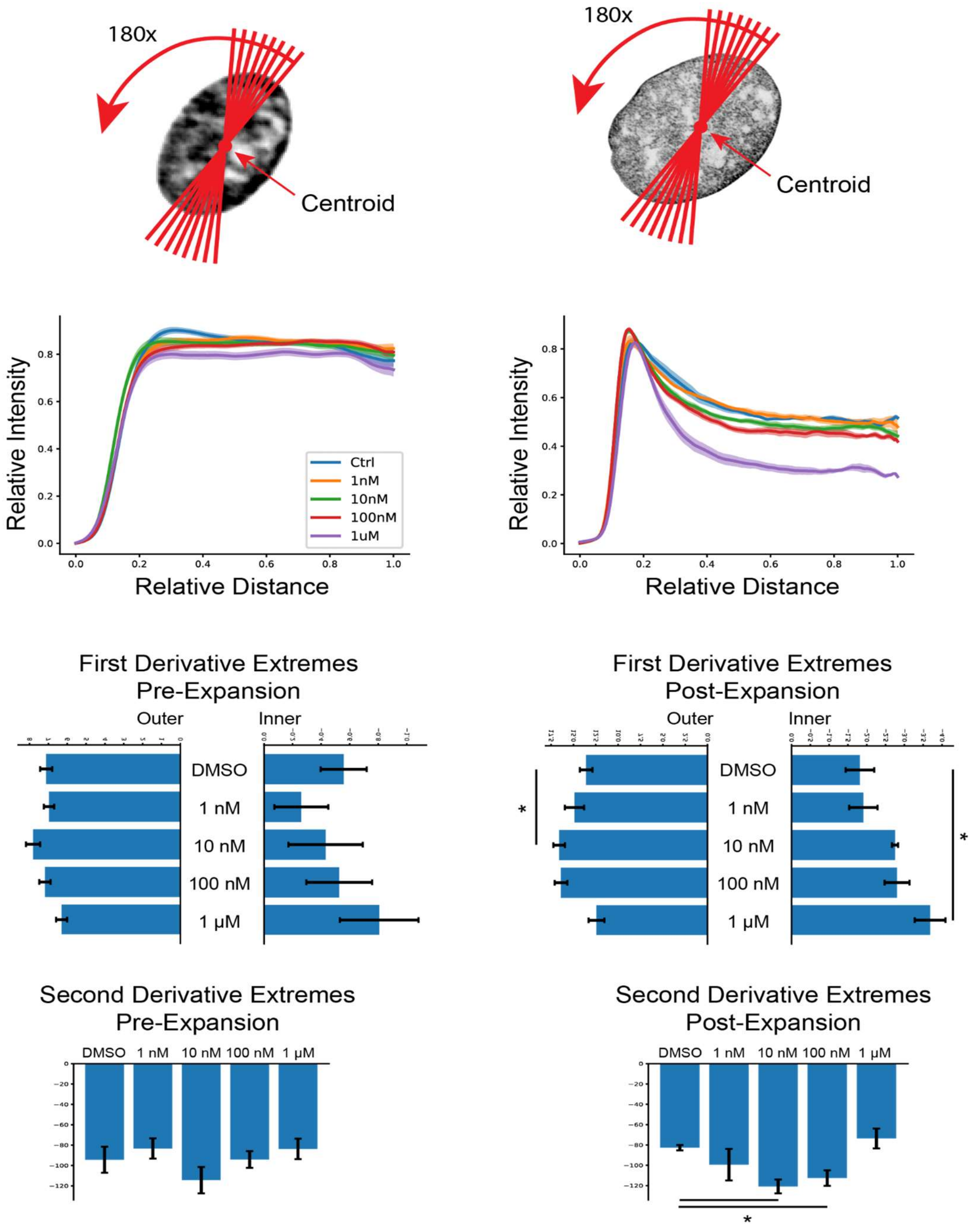


**Fig. S13: Line-scan analysis of Doxorubicin-treated cardiomyocytes.** An orthogonal line-scan analysis shows similar results in Fig. 3. Briefly, the centroid of the nucleus was found and 180 lines were drawn through the image at one-degree increments. The pixel intensities from these scans were then interpolated (linear) to normalize their lengths to 500. The scans were then cut in half, the second half of the scans were mirrored, and the two halves were averaged to give a representation of Hoechst intensity as a function of distance from the center of the nucleus. Shaded region represents SEM as in Fig. 3e. First and second derivatives were calculated as in Fig. 3g,h. Error bars represent SEM, asterisks denote significance from an independent two-sample t-test ($p < 0.05$).

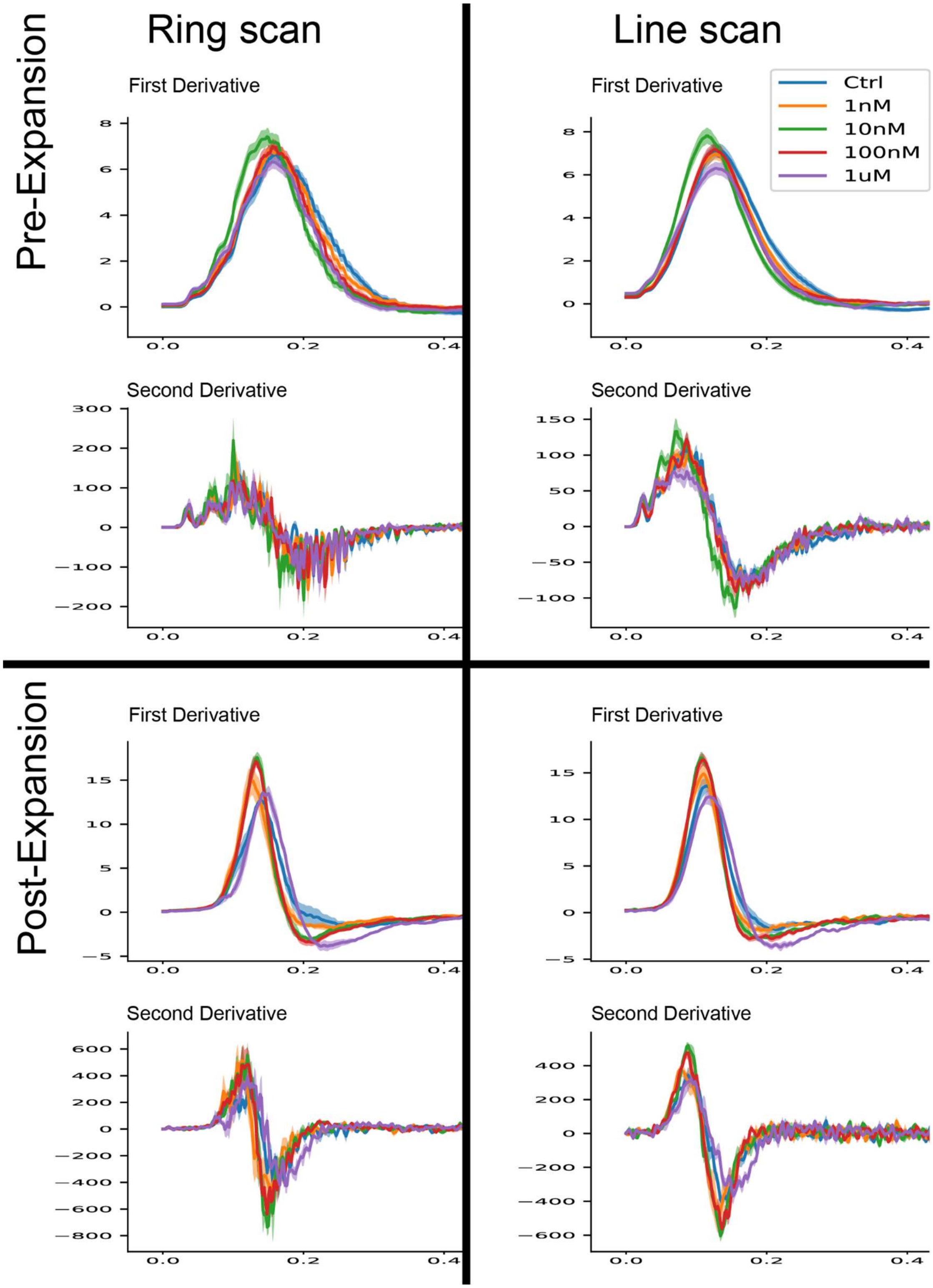


**Fig. S14: Comparison of derivative curves from Fig. 3 and Fig. S8.** Curves on the left are described in Fig. 3 (ring scan), and on the right are described in Fig. S8 (line-scan). Top 8 curves represent pre-expansion, bottom 8 curves represent post-expansion analyses.

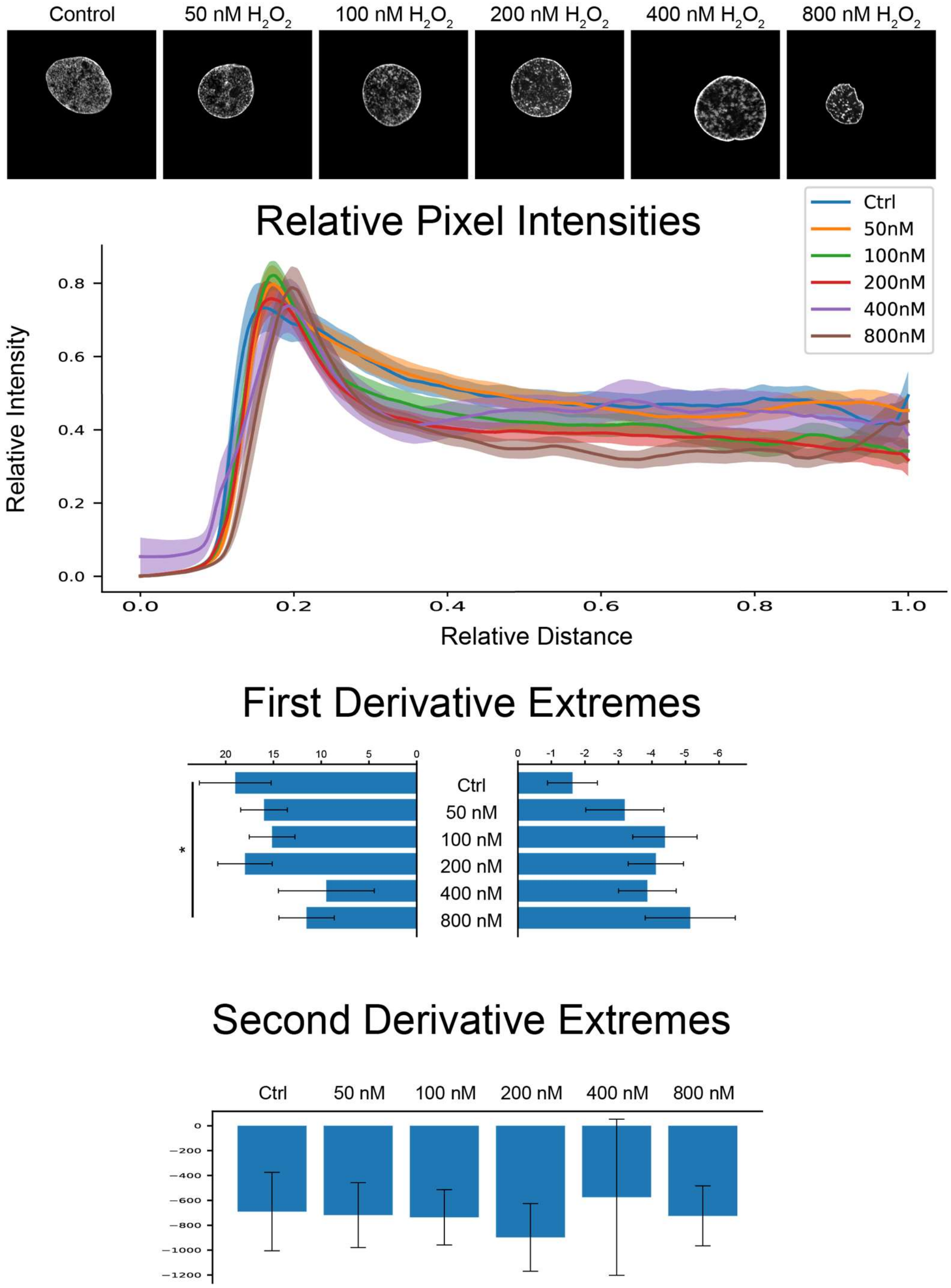


**Fig. S15: Nuclear edge analysis of $H_2O_2$-treated cardiomyocytes.** Example images of cardiomyocyte nuclei across different conditions and associated curves (top). Analyses were carried out as in Fig. 3. First derivative inflection-point-slope extremes (middle) show a significant reduction at the outer side only between the control and the highest concentration of hydrogen peroxide. Second derivative minima (bottom) show no significant differences between conditions.

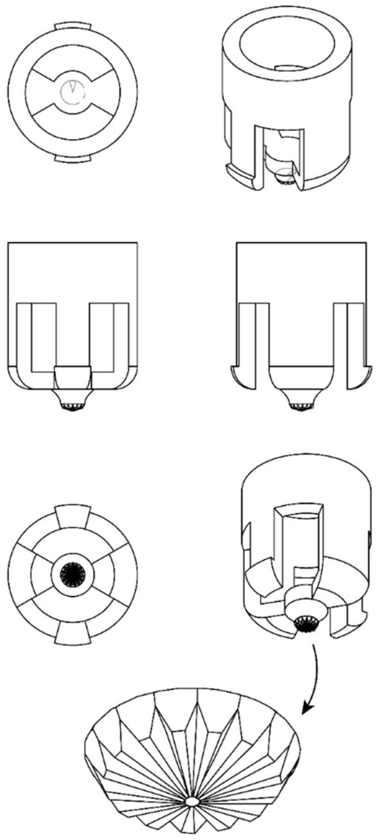

**Fig. S16: Schematic depiction of the 24-well plate HiExM device**. This device was designed to form one gel in a well of a standard 24-well cell culture plate and manufactured with injection molding (Protolabs).

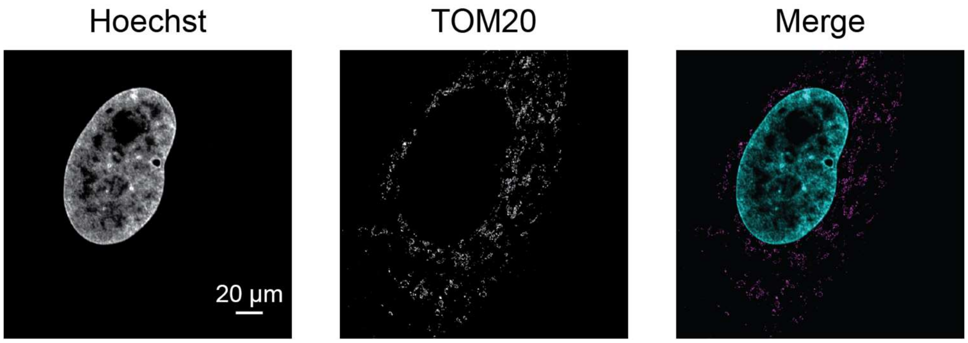


**Fig. S17: Example of a cell expanded in HiExM using Photo-ExM gel chemistry.** Photo-ExM does not require an anoxic environment for gel deposition and polymerization, improving ease of use of HiExM. Mitochondria were stained with an Alexa 647 conjugated secondary antibody, demonstrating that HiExM is compatible with additional fluorophores when combined with Photo-ExM.

### Supplementary Methods

*APS/TEMED workflow*

Fixed cells were stained using standard immunofluorescence protocols and treated with anchor solution (6-((Acryloyl)amino)hexanoic acid, abbreviated AcX) (Invitrogen A20770) for at least 3 hrs at RT prior to gelation according to standard ExM protocols.[1] Cells were then washed twice with PBS (Invitrogen AM9625) and aspirated dry. Two 2 mL vials of stock X, TEMED (Bio-Rad 1610800) and APS (Millipore Sigma A7460) (40g/100mL) were prepared and placed on ice with an aluminum block in a nitrogen glove bag (MilliporeSigma Z530112). The glove bag was purged and backfilled with dry nitrogen twice. 40 uL of APS or TEMED were added to 2 mL of stock X followed by gentle mixing. Both stock X solutions (one containing APS and the other containing TEMED) were poured onto separate absorbent pads. The first device was dipped into the TEMED-containing solution, stamped into a set of wells, and the process was repeated with the same device to load droplets of TEMED harboring stock X in all wells. Five separate devices were then dipped in the APS-containing solution, then immediately inserted into rows of the well plate. The plate was left on ice for 15 minutes, then inverted and placed in a glove bag. The underside of the well plate was covered in ~2mL of water (for thermal conduction) and ~50°C water bag was laid over the top of the inverted well plate to warm the gels for 5 min. The well plate was removed and 100 uL of digestion solution containing proteinase K (NEB P8107S) was added to each gel-containing well overnight at room temperature. The following day, the well plate was submerged in ~4 liters of DI water under constant agitation for 2 hours, exchanging the water after the first two hours. Finally, the plate was removed from the water bath and devices removed from the well plate. Gentle aspiration of remaining water from the well followed by the addition of ~100 uL of heavy mineral oil over the gel was sufficient to restrict gel movement to allow for automated imaging in a 96-well plate.

*Measurement of Volume Collected by Devices*

To estimate the volume of liquid collected by the HiExM device, the mass of water was measured on a sartorius TE64 before and after dipping the device into the reservoir. The difference in mass was measured 18 times for an average collected mass of 2806 ug +/- 482 ug. The average total mass retained by the device was divided by 12 (the number of posts per device) yielding an estimate of 234 ug or 234 nL per post.

*Ring-scan-based analysis for CM nuclei*

Analysis was identical to the analysis shown in Fig. 3 up to and including the dilation of the mask. A custom python script was used to perform the line-scan-based analysis as follows: For each nucleus, the centroid of its mask was taken and 180 lines were drawn through the center of the nucleus at one-degree intervals. For each of these lines, the intensity values for pixels through which the line passed were stored. The list of values was then split in half and the second half was flipped. The original half and the flipped half were then averaged. Finally, all 180 lines that were processed in this way were averaged to arrive at a final representation for that nucleus. Downstream steps were identical to the analysis shown in Fig. 3.

*Photo-ExM workflow*

Photo-ExM gel solutions were prepared as previously described.[2] The protocol for device use with Photo-ExM is the same as the Irgacure protocol with two exceptions. First, gel polymerization in Photo-ExM is not oxygen inhibited, so the protocol does not require a glove bag. Second, Gel solutions (2 µL) are pipetted into the wells of a custom 3D printed chip (Supplementary File PhotoExM_Gel_Reservoir.SLDPRT) designed for small volume gel solution storage in HiExM. HiExM devices are applied to the loaded chip such that each post of the device collected gel solution from one of the loaded wells.

*Fabrication of 24-well plate HiExM devices*

24-well plate HiExM devices were fabricated by Proto Labs with injection molding. The device desgin can be found using the following link:
https://github.com/lboyerlab/hiExM_Supplementary_Files

| Secondary Antibodies | Vendor | Cat. Number |
|---|---|---|
| CF350 Goat Anti-Mouse IgG (H+L) | Biotium | 20140-1 |
| CF488 Goat Anti-Mouse IgG (H+L) | Biotium | 20010-1 |
| CF543 Goat Anti-Mouse IgG (H+L) | Biotium | 20306-1 |
| CF555 Goat Anti-Mouse IgG (H+L) | Biotium | 20030-1 |
| CF568 Goat Anti-Mouse IgG (H+L) | Biotium | 20100-1 |
| CF633 Goat Anti-Mouse IgG (H+L) | Biotium | 20120-1 |

**References**

1. Asano, S. M. *et al.* Expansion Microscopy: Protocols for Imaging Proteins and RNA in Cells and Tissues. *Curr. Protoc. Cell Biol.* **80**, (2018).
2. Günay, K. A. *et al.* Photo-expansion microscopy enables super-resolution imaging of cells embedded in 3D hydrogels. *Nat. Mater.* **22**, 777–785 (2023).
