## Supplementary figures and images for "HiExM: high-throughput expansion microscopy enables scalable super-resolution imaging"

### Movie S1

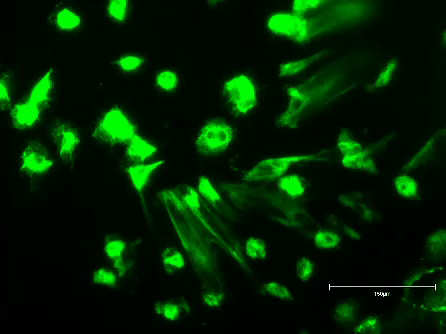

### Movie S2

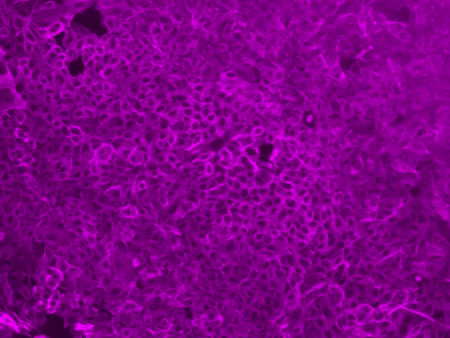

### Movie S3

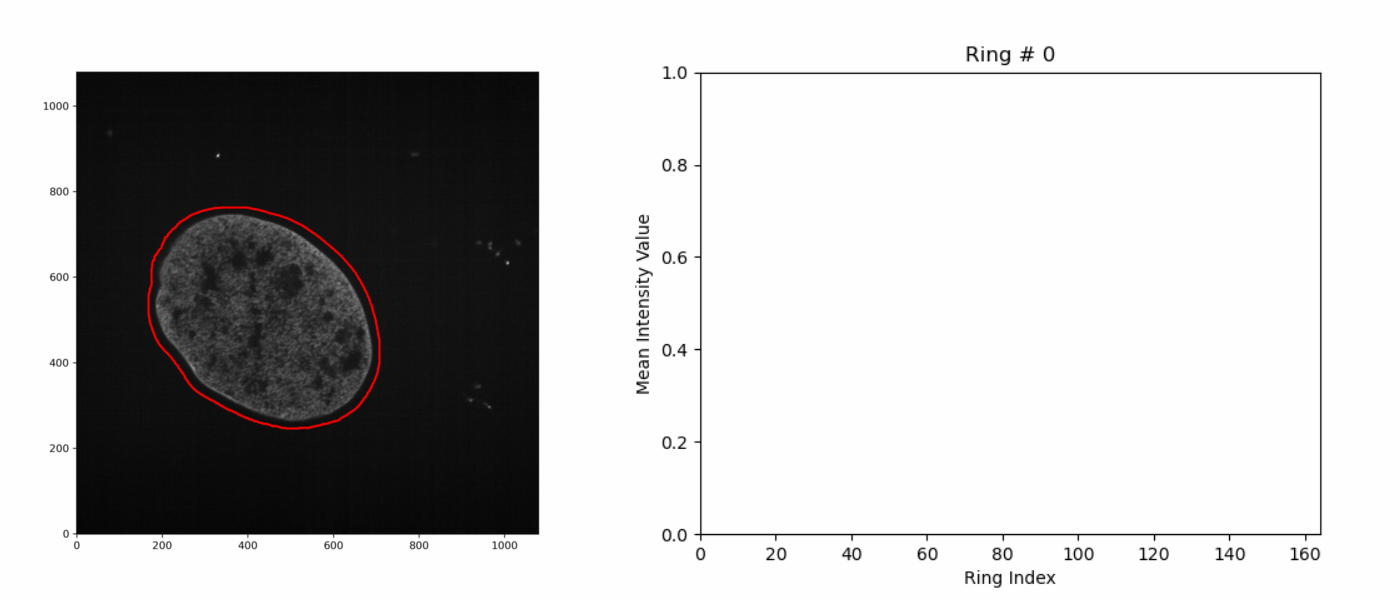
